# Principled Multi-Omic Analysis Reveals Gene Regulatory Mechanisms Of Phenotype Variation

**DOI:** 10.1101/145607

**Authors:** Casey Hanson, Junmei Cairns, Liewei Wang, Saurabh Sinha

## Abstract

Recent studies have analyzed large scale data sets of gene expression to identify genes associated with inter-individual variation in phenotypes ranging from cancer sub-types to drug sensitivity, promising new avenues of research in personalized medicine. However, gene expression data alone is limited in its ability to reveal cis-regulatory mechanisms underlying phenotypic differences. In this study, we develop a new probabilistic model, called pGENMi, that integrates multi-omics data to investigate the transcriptional regulatory mechanisms underlying inter-individual variation of a specific phenotype – that of cell line response to cytotoxic treatment. In particular, pGENMi simultaneously analyzes genotype, DNA methylation, gene expression and transcription factor (TF)-DNA binding data, along with phenotypic measurements, to identify TFs regulating the phenotype. It does so by combining statistical information about expression quantitative trait loci (eQTLs) and expression-correlated methylation marks (eQTMs) located within TF binding sites, as well as observed correlations between gene expression and phenotype variation. Application of pGENMi to data from a panel of lymphoblastoid cell lines treated with 24 drugs, in conjunction with ENCODE TF ChIP data, yielded a number of known as well as novel TF-drug associations. Experimental validations by TF knock-down confirmed 41% of the predicted and tested associations, compared to a 12% confirmation rate of tested non-associations (controls). Extensive literature survey also corroborated 62% of the predicted associations above a stringent threshold. Moreover, associations predicted only when combining eQTL and eQTM data showed higher precision compared to an eQTL-only or eQTM-only analysis with the same method, further demonstrating the value of multi-omic integrative analysis.

## Introduction

There is great interest today in understanding why certain drugs are effective in some individuals but less so in others. A panoply of studies has sought to identify mechanisms of action of specific drugs (1) (2) (3) (4) (5) as well as genotypic variations that are predictive of an individual’s drug response (6) (7) (8). A major class of drugs of interest today are cytotoxic drugs that may be used in cancer treatment. Large scale data generation efforts, including genotypic and molecular profiling of panels of cell lines (9) (10) along with drug response (cytotoxicity) measurement on those cell lines (9) (11) (12), are expected to facilitate future advances in cancer pharmacogenomics. As the diversity of such data sets increases, it is important to devise rigorous computational methods that can combine these diverse data in a principled manner to help scientists answer mechanistic as well as therapeutic questions pertaining to drug response. For instance, correlating gene expression and drug response in a panel of cell lines helps identify cytotoxicity-related genes (12) but it is not clear how one might extend the approach to additionally exploit genotype (SNP) and epigenotype data (e.g., CpG methylation marks) on the same cell lines to maximum effect. We need a statistically sound approach capable of modeling phenotypic variation while integrating several heterogeneous genomic and epigenomic data types.

One of the key mechanistic questions related to drug response variation, or indeed for any phenotypic variation under study, is the role of gene regulatory networks (GRNs) in shaping such variation. GRNs are a natural formalism for describing global gene expression dynamics and variation thereof, in a succinct and interpretable manner (13) (14) (15). A major step in characterizing GRNs mediating drug response variation is to identify functionally important transcription factors (TFs), as TFs are the main actors in any GRN (16) (17) (18). Identified TFs may then be experimentally validated by showing that their knock-down leads to change in chemosensitivity (19) (20) (21). Mechanistically speaking, a TF may influence drug response by regulating one or more target genes whose expression levels are in turn linked to the strength of cytotoxic response, e.g., if the target genes are involved in apoptotic pathways. In such a case, one expects evidence of the TF’s regulatory influence on the target gene(s) in the form of binding sites revealed as ChIP peaks (19).Thus, if one finds substantial evidence that TF binding sites harbor genotype or epigenotype variations that are in turn correlated with gene expression and to the phenotype, it should be possible to statistically implicate that TF in drug response variation. This is the key insight pursued in this work, to identify major transcriptional regulators of drug response variation across individuals.

There have been a number of studies linking specific phenotypes such as disease status to elements of the non-coding genome (22) (23) (24) (25), many utilizing the NGHRI Genome Wide Association (GWA) Catalog SNPs (26). These studies have been facilitated by large scale efforts such as the ENCODE Project (27) and the Epigenomics Roadmap Project (28) that provide a guide to identifying non-coding elements of the genome, which can then be used as a regulatory context to interpret GWA SNPs. Additionally, there have been studies that quantify the impact of non-coding genetic variation on molecular profiles such as TF-DNA binding or DNA accessibility (29) (30) (31) (32) (33), often utilizing the NHGRI GWA catalog to assess the phenotypic consequences of variants. Despite numerous efforts to connect phenotype with genotype and regulatory elements, there has not been a systematic effort to aggregate such connections to learn major regulatory mechanisms underlying the genotype-phenotype relationship and its variation across individuals. Furthermore, the regulatory impact of epigenetic sources of variation (for instance CpG methylation) are usually assessed in isolation of genetic variants (SNPs). Recent studies have shown that there may be a complex interplay between genetic, epigenetic, and transcriptional variation in relation to disease (34), arguing for a more integrative approach to their analysis.

We present here a novel, statistically principled approach to aggregating data on genetic as well as epigenetic variations, along with genome-wide profiles of regulatory function, to derive associations between a transcription factor and individual variation in cytotoxic response to a drug; this permits a mechanistic interpretation of the impact of molecular variants on drug response. There have been new insights into how TFs may be regulated by small molecules and other mechanisms such as modulating protein-protein interactions (35) (36) (37) (38), and there is no doubt as to their significance in human diseases such as cancer (39) (36) (40) (41) (42). Our aim therefore is to push the frontier of knowledge regarding the relationship of TFs and drug response for putative therapeutic benefit. Our new computational method is based on a statistical formalism called probabilistic graphical models (43), which are among the most flexible and principled ways available today for inference from heterogeneous and noisy data. It builds on a previous approach, Gene Expression in the Middle Analysis (or GENMi), that relied on enrichment tests and is sensitive to arbitrary thresholds (19). Using a rigorous data preprocessing pipeline in conjunction with this powerful framework, we demonstrate that the TF-drug associations predicted by the method are accurate, by showing that knock-down of the TF affects sensitivity to the drug. The new method is also applicable to other studies where one seeks mechanistic factors underlying individual variation in a quantitative phenotype in the presence of genotype and epigenotype information. We present our results in the first online compendium dedicated to TF-drug associations confirmed experimentally, available at the following location: veda.cs.uiuc.edu/pgenmi

## Results

### A new probabilistic model to integrate genotype, gene expression and phenotype data

In our previous work, we presented a proof-of-principle method that identifies TFs associated with drug response (or any other quantitative phenotype) by integrating over multiple data types (19). We searched for cases where a SNP in the cis-regulatory region of a gene is correlated with the gene’s expression (cis-eQTL analysis) and the gene’s expression is correlated with phenotype (44); if significantly many cases like this were identified involving SNPs within ChIP peaks of a TF, then the TF was deemed to be associated with individual variation in phenotype. In other words, SNP – gene expression – phenotype relationships were discovered first and the identified non-coding SNPs were examined for evidence of a common mechanistic theme – their location within a TF’s binding sites. Here, we constructed a rigorous probabilistic model that builds upon this basic principle to identify phenotype-related TFs. The new method is called ‘pGENMi’ (the previous tool was named ‘GENMi’ for ‘gene expression in the middle’ and the ‘p’ denotes a probabilistic model). It integrates information from many genes whose expression correlates with the phenotype, and where for each gene it can find one or more types of evidence supporting a regulatory role for a specific TF. The probabilistic formulation of pGENMi offers the following important features:

1. It integrates expression-phenotype associations (TWAS) without relying on strict thresholds.
2. To quantify evidence for a TF’s role in gene expression variation, it can utilize information from cis-eQTL SNPs as well as other types of variants, including CpG methylation and other epigenetic variants, located in the TF’s binding sites near the gene.
3. In incorporating multiple sources of evidence for a TF’s regulatory role, it is able to weight the contribution of each type of evidence differently, learning these relative weights automatically.

The new method computes a p-value for the correlation between the phenotype and a gene’s expression (TWAS), and models this p-value using a mixture of two distributions: a null (uniform) distribution, and a beta distribution that is biased towards significant p-values. These two component distributions reflect the cases where the gene is or is not ‘associated’ with the phenotype, a relationship included as a hidden binary variable in the model. The existence of this gene-phenotype relationship is, in turn, modeled as a probabilistic function of multiple lines of evidence supporting a TF’s role in regulating the gene. The model can accommodate an arbitrary number of regulatory evidences, such as the existence of a significant cis-eQTL within the TF’s binding site located near the gene. Specifically, the probability of the gene being associated with the phenotype is modeled as a logistic function of a linear combination of evidences, using a single free parameter for each type of evidence representing the contribution of that type towards the gene-phenotype relationship (the free parameter is shared across all genes, making it well constrained). Finally, pGENMi provides a log likelihood ratio score to quantify a TFs’ roles in regulating the phenotype in light of all available data. We describe the probabilistic graphical model of pGENMi in detail in **Methods** and in **Figure 1**.

**Figure 1:**
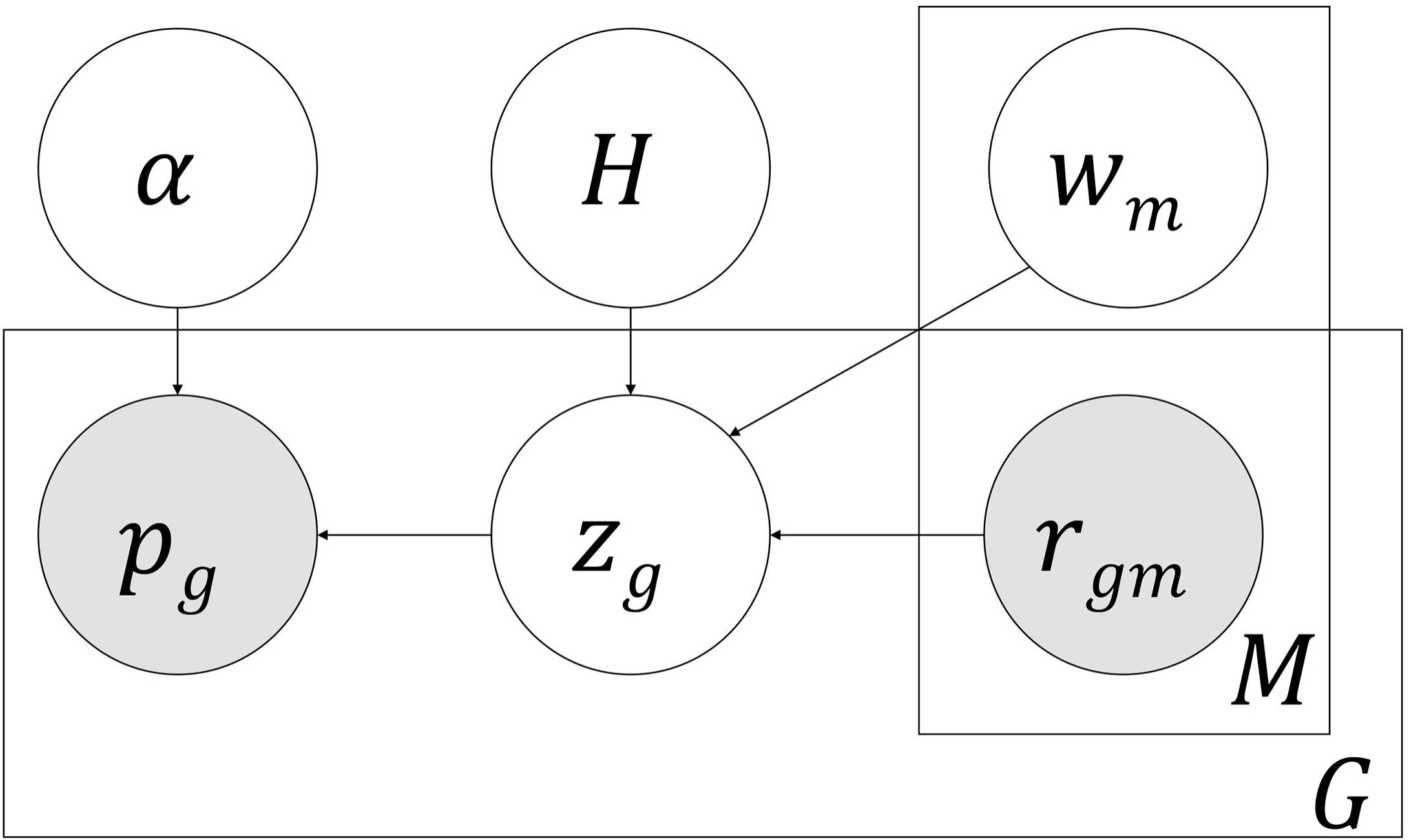
Plate diagram of pGENMi model.

### pGENMi integrates multiple types of regulatory evidence to elucidate the role of TFs in drug response variation

We applied the pGENMi algorithm to identify TFs that putatively regulate individual variation in drug response. We analyzed drug response data of 24 cytotoxic drugs (or treatments) separately. Genotype (SNP), methylation, and gene expression data (using a principal component gene expression representation derived from probe data, see **Methods**) on 284 lymphoblastoid cell lines (LCLs) were analyzed in conjunction with cytotoxicity data for the same panel of LCLs, as well as ENCODE ChIP-Seq data on TF-DNA binding. To assess the regulatory evidence for a TF’s role in a gene’s expression variation, we first identified the strongest cis-eQTL SNP located near the gene (50Kb upstream of the transcription start site) and within the TF’s binding sites. We applied a threshold of 0.05 to the eQTL p-value to designate the presence of eQTL evidence for the TF’s role in regulating the gene. We adopted the same approach to designate the presence of eQTM evidence (the correlation of methylation marks with gene expression, see **Methods**) and restricted our analysis to methylation marks near the gene and within TF binding sites. It should be noted that for every possible correlation analysis (TWAS, eQTL, and eQTM), we included as covariates all information for which we had data per individual (sex, age, batch) as well as data we could infer from ethnic labels and genotype information (sub-population axes of variation from EIGENSTRAT (45)) – see **Methods**. The eQTL and eQTM evidence (or lack thereof) for the TF’s regulatory influence on genes was then integrated with gene expression-phenotype associations (TWAS) by pGENMi. The result of this analysis, which we refer to as eQTL+M, is a ranking (by LLR score) of all TFs by their predicted role in regulating each drug response phenotype. We also repeated the entire procedure using eQTL-only evidence and eQTM-only evidence.

### Literature evidence in support of pGENMi predicted (TF, Treatment) associations

In the absence of a gold standard dataset, we performed an extensive literature search to find publications with experimental results corroborating our reported associations. When scanning the literature, we considered whether a study validated a reported (TF, Treatment) pair using two different criteria. The first, and most convincing, is experimental evidence demonstrating that knockdown of the TF affects chemosensitivity of a cell to the corresponding treatment; we call such evidence “direct”. The second is experimental evidence demonstrating differential expression or DNA binding of the TF induced by the treatment; we consider such evidence to be valuable, but less convincing than “direct” evidence and consequently label it as “indirect”. To begin our evaluation, we restricted our literature search to eQTL+M associations with LLR ≥ 4.5, roughly corresponding to a p-value of 0.05 (χ^2^ test). Of the 19 reported associations at this stringent criterion, 14 were validated by the literature (8 directly and 6 indirectly), as shown in **Table 1** and presented as a detailed survey in **Supplementary Text 1.** As an example, pGENMi reported *FOXM1* as a top scoring association for the drug Temozolomide; this prediction was validated by a study where siRNA knockdown of FOXM1 was shown to sensitize recurrent glioblastoma multiforme (GBM) tumors to this drug (46). pGENMi also reported *STAT3* as a top candidate TF associated with response to the drug oxaliplatin. Indeed, siRNA silencing of STAT3 combined with oxaliplatin therapy, in mouse models of metastatic colorectal cancer (HCT116), was previously found to reduce tumor size by 96%, better than either treatment separately (77% and 57% respectively) (47). An example of indirect validation is that of the predicted association between *GATA1* and rapamycin. Treatment of hexamethylenebisacetamide (HMBA), which commit cells to cessation of growth and differentiation, to friend erythroleukaemia cells increased DNA binding of GATA1, an important transcription factor for erythroid specific genes. When treated with the S6-kinase inhibitor, rapamycin, HMBA cells induced at 18 hours showed markedly lower binding of GATA1 to the DNA (48).

Some drug responses may be easier to model using gene expression than others, due to variation in the quality of experiments and different mechanisms of action. As such, we sought to examine how literature-based validations of pGENMi associations decomposed by class of drug. To do this, we relaxed the LLR threshold to 3, but restricted the search to the top seven TF associations for each drug. Figure 2 illustrates these 90 (TF, Treatment) associations in a heatmap colored by LLR and **Table 2** shows the precision (fraction of positive predictions that are validated by literature evidence) of the top associations for each drug. The results vary widely across drug classes. For certain drugs, such as cisplatin, all seven predictions were validated, either through direct or indirect literature evidence. We also observed 100% precision for the following treatments for which a single TF was predicted: arsenic, hypoxia, and docetaxel. On the other hand, TFs associated with certain treatments, e.g., 6-MP and 6-TG, were rarely supported by the literature. Overall, our literature survey (detailed in **Supplementary Text 2**) resulted in validation of 45 of 73 (62%) predictions made at the modest threshold of LLR >= 3, but pointed to substantial inter-drug variability in the ability of our method to identify TF determinants of cytotoxicity.

**Figure 2:**
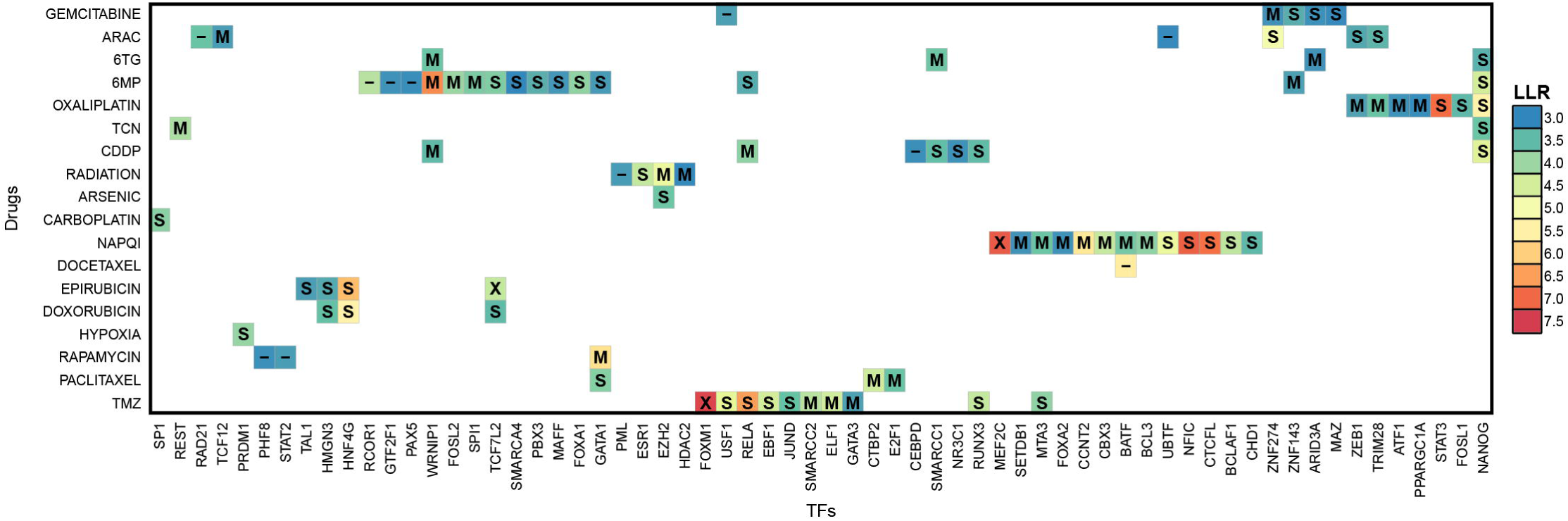
The 90 (TF, Treatment) associations predicted by eQTL+M with LLR >= 3, colored by LLR. Labels indicate whether eQTL or eQTM analysis corroborated the prediction at an LLR >= 1.74. A “X” indicates both eQTL and eQTM analysis supported the prediction, while a “S” indicates eQTL-only analysis supported the prediction. A “M” indicates eQTM-only analysis supported the prediction, while a “-“ indicates the prediction is unique to eQTL+M.

We also assessed the benefit of modeling multiple types of regulatory evidence simultaneously (i.e., eQTL+M) rather than simply taking the union of associations reported by running pGENMi for each evidence (eQTL or eQTM) separately. To answer this question, we ran eQTL and eQTM models and applied an LLR threshold of 1.74 (roughly corresponding to *χ*^2^ p-value of 0.05) for their reported associations. We then categorized the 90 eQTL+M associations with LLR ≥ 3 based on their recapitulation in eQTL- and eQTM-only analysis, as shown in **Table 3**. Strikingly, the eQTL+M associations were rarely supported by both analysis. Despite this, 11 associations were identified by the eQTL+M model that a simple intersection would not produce; we further investigated these associations in our experimental validation. In looking at the top 500 (TF, drug) associations reported by each analysis, we noted that the eQTL+M analysis and eQTL-only analysis showed strong concordance (309 common associations), while the eQTM-only analysis showed much less concordance with results from the other two analyses **(Supplementary Tables 1–3).**

### Experimental validation

We sought to experimentally verify whether TFs associated with drug response variation (by pGENMi) could be linked *in vivo* to significant changes in cellular sensitivity to drug-induced cytotoxicity. We selected predicted (TF, Treatment) pairs that reflect a diversity of regulatory support, shown in **Table 4A** and **4B**, for which we did not find prior literature support. For instance, we selected 10 of the 11 (TF, Treatment) pairs identified uniquely by eQTL+M analysis (we omitted *GTF2F1)*, and two of the three (TF, Treatment) pairs predicted by all three analyses: eQTL+M, eQTL, and eQTM. Additionally, we selected the top four pairs by LLR in the eQTL+M analysis, that were also supported by eQTM analysis. We also included a high-scoring association from eQTM-only analysis that scored poorly in the eQTL+M analysis (i.e., LLR < 3). All 17 of the these (TF, Treatment) pairs were reported by pGENMi models consistent with our regulatory hypothesis, i.e., where presence of regulatory evidence makes the gene more likely to be associated with drug (parameters w_i_ > 0, for i ≠ 0, see Methods). Some associations did not obey this strict criterion, but received high LLR scores from pGENMi. We included eight such (TF, Treatment) associations, as a strong, negative control group, pushing the total number of experimental pairs to 25. Note that we expected these eight predictions to prove false, since pGENMi considered them as statistically interesting but not in a manner consistent with its mechanistic underpinnings. In selecting these 25 candidates for experiments, we avoided TFs like *BRF2* and *GTF2F1* that were associated with 10 or more drugs. These are general transcription factors whose association is likely due to having many more ChIP peaks than other TFs, a point we have discussed in previous work (19). We also excluded NAPQI, a toxic byproduct produced during the xenobiotic metabolism of the analgesic paracetamol, due to its lack of clinical application.

Though we utilized lymphoblastoid cell lines data for our association analysis, we performed siRNA knockdown experiments in several different cell lines to demonstrate the generalizability of our results and the LCL model system. Based on clinical relevance, the human triple negative breast cancer MDA-MB-231 cells were chosen to test anthracycline, taxane, platinum, gemcitabine, radiation, and rapamycin mechanisms of action, because these drugs are typically administered as first-line therapy for triple negative breast cancer. We used a human leukemia Jurkat cell line to test 6-MP, 6-TG, and Ara-C since these several drugs are used to treat leukemia. Temozolomide is the first line therapy for glioblastoma multiforme; we therefore used human glioma U251 cells to validate the TFs associated with Temozolomide response. The siRNA knockdowns were performed for the 25 (TF, treatment) pairs shown in **Table 4A** and **4B** for the eQTL+M predicted and negative control associations respectively. Cytotoxicity graphs of all knockdowns are shown in Figure 3. Overall, 7 of 17 (~41%) of predicted associations, shown in **Table 4A**, were validated in this way, compared to 1 of 8 (~12%) of negative control associations, shown in **Table 4B**. Additionally, the precision on the 10 unique eQTL+M predicted associations was 50% (5 of 10), compared to 0% (0 of 2) of eQTL+M negative control associations; this indicates pGENMi is able to combine multiple lines of regulatory evidence to make novel regulatory predictions with reasonable precision that would otherwise be unreported when considering each line of evidence in isolation. With respect to the single (TF, Treatment) pair predicted uniquely from eQTM analysis (and not predicted by the other two modes of analysis), we failed to validate it in our experiments. Nevertheless, we believe that our experimental validation demonstrates the overall utility of pGENMi predictions overall and especially when combining multiple sources of regulatory information to learn novel associations.

**Figure 3:**
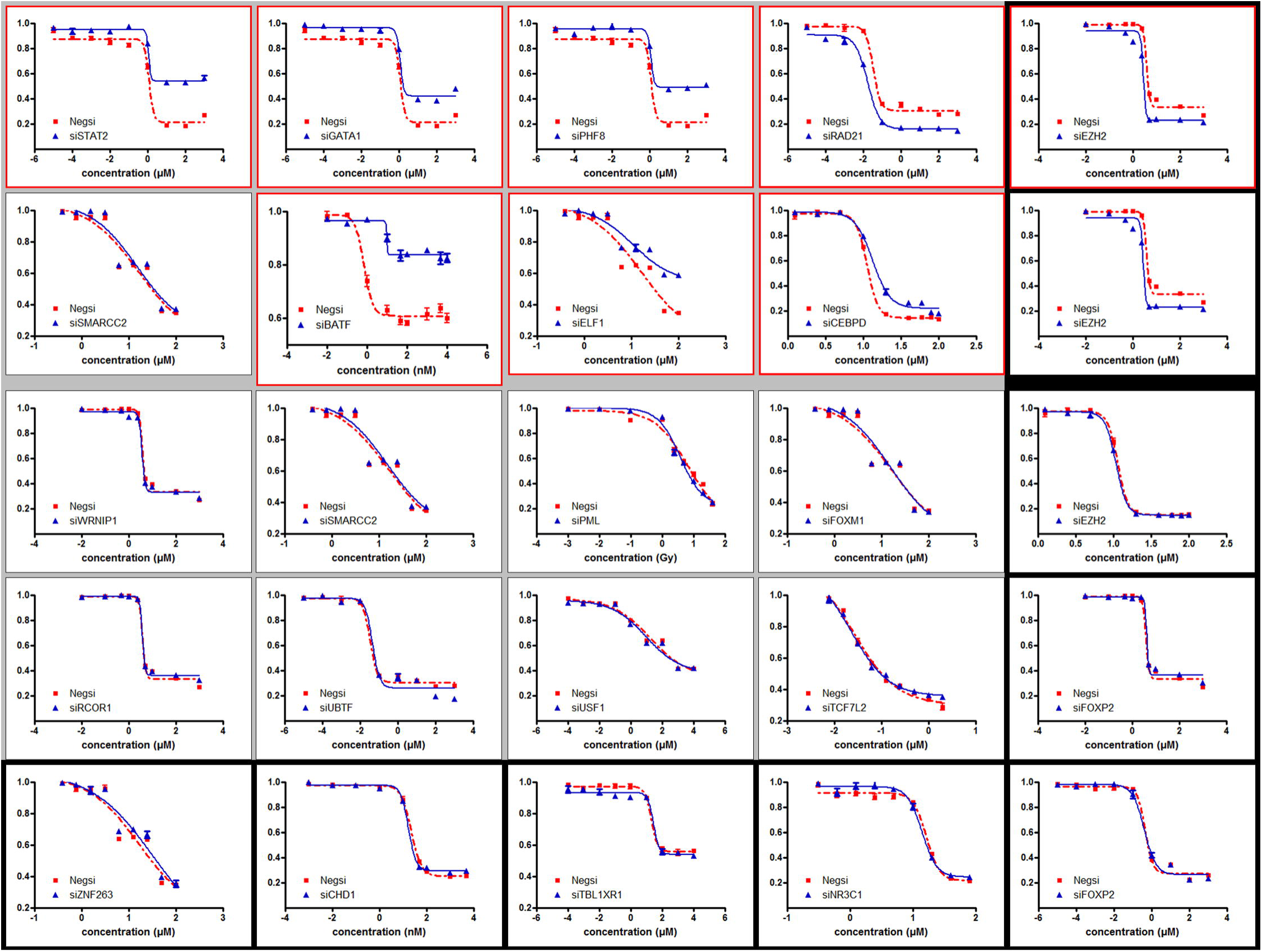
All 25 experimentally tested (TF, Treatment) curves. Red outlines indicate significant shifts in cytotoxicity between siRNA negative and siRNA TF conditions. Curves with a gray background are eQTL+M predictions, while those with a black background are negative controls. We validated 7 out of 17 predictions consistent with our regulatory hypothesis and only 1 out of eight negative controls.

### pGENMi produces more associations than a naïve baseline

The pGENMi algorithm, with results restricted to those with an LLR >=3, produced 90 (TF, Treatment) associations. As no such algorithm other than GENMi (19) exists for associating TFs with drug response, and even GENMi is incapable of handling both genotype and methylation information simultaneously, there was no choice of baseline method to compare the results to. However, we did compare pGENMi to a simple TWAS correlation approach, where a TF is considered associated with a drug if the TF’s expression (mRNA level) correlates significantly with cytotoxicity. This approach has been recently proposed for identifying genes associated with drug response (12). After FDR correction across all 148 TFs for each drug separately at the 0.10 threshold, this baseline approach reported 103 associations. However, these associations were mostly distinct from those reported by pGENMi: between the 103 associations reported by the TWAS method and the 90 associations reported by pGENMi, only the following five associations are shared: (Carboplatin, SP1), (6-MP, TCF7L2), (6-MP, SMARCA4), (6-TG, SMARCC1), (Cisplatin, SMARCC1). This demonstrates that pGENMi is able to reveal novel insights into regulatory mechanisms of drug response, based on cis-regulatory analysis, that may not be obtained from the current approach of expression-phenotype correlations.

### Database of (TF, Treatment) Associations

We have made available as an online resource all (TF, Treatment) associations validated using siRNA or overexpression experiments in this study as well as those found to be similarly validated in our survey of the literature (**Supplementary Text 1 and 2**), the GENMi study (19), and a related work that performs the same experimental validations (49), This resource is available at veda.cs.uiuc.edu/pgenmi. We believe this to be the first such catalog of experimentally validated (TF, Treatment) pairs.

### Discussion

In this work, we developed a principled probabilistic graphical model for inferring TFs that regulate drug response variation among individuals. To our knowledge no prior study aims to answer this question to which we can compare our methodology. Nevertheless, we have demonstrated a high rate of success in validating our associations through a comprehensive literature survey and *in vivo* experimental validation. We have also collated these successful validations from literature and the lab along with the results of various studies into the first (TF, Treatment) association database.

While we view the pGENMi model as an major improvement over our previous proof of principle approach (19), it is similar to various existing models for data integration and may be further improved upon in many ways, from model to experimental design. pGENMi is similar to the GPA model (50), except we do not model SNPs as latent variables (instead genes), and use an empirical Bayesian approach similarly used to prioritize SNPs using annotations (33) and Battle et. al’s latent regulatory variant model (LRVM) (51, 52). Additionally, our model is also similar to EPS (53), except we do not model gene expression directly, but rather regulatory information, and use a logistic instead of a Gaussian distribution. pGENMi can be improved in a number of ways; for instance, our model makes the implicit assumption that a latent variable *z_g_* representing a gene’s involvement in mediating the TF’s influence on phenotype is independent across genes; however, co-regulated gene sets break this assumption and it would be more reflective of the underlying biology to model a network of latent variables *z_g_* instead of treating each gene independently. Our choice of modeling the p-values of gene expression-phenotype correlation as arising from a Beta distribution avoids the use of arbitrary thresholds on the strength of this relationship, but we believe there may be less restrictive ways to explicitly model phenotype-expression relationships. Finally, the pGENMi does not model the full signal transduction pathway from membrane receptors to nuclear proteins; utilizing phosphorylation data has already been utilized to infer context specific casual signaling (54) and could empower us to identify those TFs likely to be stimulated by the drug. However, taking a more systemic view would require expensive changes to the model and is left for future work. Although we integrated more types of data in this work than typical studies that directly relate gene expression to drug response data (12) (52), the model allows further extensions. We restricted ourselves to eQTL and eQTM data (in addition to gene expression, phenotype, and TF-DNA binding data) because that was the extent to which we had variant profiling on the LCLs. Incorporating the impact of various histone modifications and other epigenetic marks would have been an interesting endeavor.

Additionally, it may be wise to consider carefully designing *in vivo* assays to validate the mediators of drug response, rather than the TFs themselves, in the future – even if it means decreasing the number of associations that can be validated. Despite these various areas of improvement, the pGENMi methodology was successful in integrating both genetic and epigenetic variation to elucidate the regulatory association between a TF and treatment, as evidenced by our 50% precision in validating those predictions only eQTL+M reported. We hope this methodology can serve as a blueprint for future endeavors that aim to elucidate the regulatory basis of disease etiology using multiple molecular profiles of variation.

The pGENMi framework need not be restricted to the analysis of the variation of drug response. It may be easily adapted for use in other situations where one seeks to link a phenotype to set of gene properties, e.g., regulation by a TF, membership in a pathway, involvement in a biological process, etc., while using gene expression as an intermediate variable. The relationship between gene expression and phenotype may be quantified by differential expression p-values if, for instance, the phenotype is binary as in case vs. control studies or in before vs. after treatment studies. The method tries to find the most likely mixture model where the latent assignment of the gene expression-phenotype association to a model (in this case, null or alternative) is distributed according to a logistic function of observed properties of the gene. In this work, the observed properties reflect TF regulation and the phenotype is drug response; thus, we are able to relate drug response to transcription factors via multiple sources of regulatory evidence.

## Methods

### Data Acquisition

We sequenced single nucleotide polymorphisms (SNPs), 3’ mRNA probe expression, and CpG methylation status across 284 Epstein-Barr Virus (EBV) transformed lymphoblastoid cell lines (LCL) from the Coriell Cell Repository. The ethnicities of the cohort decomposed along the following three broad ethnic lines: 95 Han-Chinese American (HCA), 96 Caucasian American (CA), and 93 African American (AA). The sequenced genotype data resulted in 1,362,404 germ line single nucleotide polymorphisms (SNPs), each with minor allele frequency (MAF) < 5%, genotype rate > 95%, and in Hardy-Weinberg Equilibrium. Imputation analysis of this initial set of genotyped SNPs resulted in 11,256,504 SNPs. Gene expression data consisted of 54,613 Affymetrix U133 Plus 2.0 Gene-Chip probes, transformed using log_2_ GC Robust Multi-Array Averaging (GC-RMA). Genotyped SNP and gene expression data are available in the NCBI Gene Expression Omnibus (GEO) (http://www.ncbi.nlm.nih.gov/geo) under SuperSeries accession no. GSE24277 and were originally published in a study by Niu et. al (55). In addition to gene expression and genotype data, our collaborators at the Mayo Clinic sequenced 444,797 methylation marks, originally published by Heyn et al. (56), and made available in the NCBI GEO under SuperSeries accession no. GSE36369. As a representation of methylation status, we used the “beta” value, which encodes methylation status within [0, 1], where 0 and 1 correspond to total absence and presence of the mark respectively.

Drug response data was derived from dosage-response curves of 24 cytotoxic treatments: 6-MP, 6-TG, Ara-C, arsenic, carboplatin, cisplatin, cladribine, docetaxel, doxorubicin, epirubicin, everolimus, fludarabine, gemcitabine, hypoxia, metformin, MPA, MTX, NAPQI, oxaliplatin, paclitaxel, radiation, rapamycin, triciribine, and temozolomide. Each response curve was summarized by an EC50 value (drug concentration at which half the original LCL population survived treatment). These data were most recently analyzed by Hanson et al. (19). Experimental data on TF binding were obtained from the ENCODE project (57). The union of Clustered ChIP (v3) tracks of 161 TFs across 91 cell lines formed a single track for each TF, representing a TF composite regulatory profile reflective of activity across a wide variety of cellular contexts. This follows the methodology employed by (19). Contrary to that paper, we exempted the following 13 TFs from the full 161, leaving 148 for analysis: general TFs *(POLR2A, POLR3A, POLR3 G, TBP)* and those for which no eQTL or eQTM SNPs (p < 0.05) were detected within ChIP peaks *(BDP1, BRCA1, BRF1, ELK1, ELK4, ESRRA, HSF1, KDM5A, NELFE).*

### Pre-Processing of Data

#### SNP imputation

SNP imputation was performed as in (19), using the BEAGLE v3.3.1 software (58) (See **Supplementary Text 3** for details).

#### Summarizing probe expression by gene symbol

Rather than modeling raw probe data, we opted to summarize probe level expression at the Ensembl gene symbol level, since representing gene expression using multiple correlated probes breaks the independence assumptions of our probabilistic model by giving greater weight to genes with higher probe coverage. Of the 54,613 probes, those with low variance (*σ*^2^ ≤ 0.1), were omitted from analysis, purging 5,408 probes. Of the remainder, 34,832 probes mapped to 16,183 stable Ensembl gene symbols with at least one exon annotation, using the GRCh37 Ensembl BioMart mapping of Ensembl gene symbols with HGU 133 Plus 2.0 array probe identifiers. Since multiple probes can map to the same gene, we utilized a simple algorithm to obtain a single gene expression value as a function of its mapped probe expressions. The following illustrates this procedure, which uses principle component analysis (PCA) to obtain a vector of gene expression and Z-score normalization to ensure all genes are scaled identically:

For each gene *g* with at least one mapped probe:

1. If *g* has one mapped probe

a. Return z-score normalization of the probe’s expression.
2. If gene *g* has *n* mapped probes with a probe expression matrix of size *n* × 284:

a. Obtain the first principal component (PC_1_) of the probe expression matrix.
b. Project the probe expression matrix onto PC_1_.
c. Return z-score normalization of the projected PC1 expression.

Applying this procedure resulted in a matrix of Ensembl gene expression with dimensionality 16,183 (number of genes) × 284 (number of LCLs), and Z-score normalized expression across individuals. Henceforth in this publication, we refer to this matrix and its values as the gene expression matrix and gene expression, respectively.

#### Controlling for confounding variables

We included the following potentially confounding variables as covariates in all regression analyses for this study: batch, age, gender, and sub-population labels, derived from EIGENSTRAT (see below). We decided to omit explicit labels of ethnicity, as the sub-population labels should capture that information implicitly. To derive p-values for the covariate of interest in the multiple regression (for instance, a SNP in an eQTL regression) we computed the p-value of the log likelihood ratio between models with and without the covariate of interest, using a *≥*^2^ distribution.

#### Population stratification

We utilized the EIGENSTRAT program (version 6.0.1) developed by the Price lab (45) (https://www.hsph.harvard.edu/alkes-price/software) to identify sub-population labels from genotype data and remove their potential confounding effects in regression analyses (see **Supplementary Text 3** for details.) We first computed a set of independent SNPs using the PLINK program (1.90 beta) (59). We then estimated sub-population labels as continuous axes of variation by using EIGENSTRAT to perform PCA on each ethnic group’s independent SNP genotype matrix, retrieved all PCs with Tracy-Widom p-values ≤ 0.05, resulting in a total of seven axes, shown in **Supplementary Table 2**. We projected the entire genotype matrix onto each of the seven axes (setting to 0 SNPs not contributing to the axis) to derive numeric sub-population representations for the cohort.

#### TWAS, eQTL, and eQTM analysis

Each of the following regression analyses controlled for the following covariates: sex, age, batch, and population axes of variation derived from EIGENSTRAT. Transcript wide association analysis, or TWAS, (44) and eQTL analysis were performed following (19). To perform TWAS, we computed Pearson correlation coefficient p-values between gene expression and drug EC50 values across the 284 LCLs, for each (gene, drug) pair. For eQTL analysis, we operationally defined the 50Kb upstream region of each gene as its putative cis-regulatory region. We performed eQTL analysis for each gene locus by calculating Pearson correlation p-values between each gene’s expression and the genotype (measured by allelic dosing of 0,1,2) of each SNP in its cis-regulatory region, provided the SNP’s location was co-incident within a ChIP peak of the TF of interest. To avoid the statistical artifacts of linkage disequilibrium, we preserved only the most significant eQTL-SNP, by correlation p-value, in each gene’s cis-regulatory region. To Principled Multi-Omic Analysis Reveals Gene Regulatory Mechanisms Of Phenotype Variation | 20 incorporate methylation as a source of epigenetic variation that could impact the binding behavior of TFs, we repeated the eQTL analysis while substituting SNP genotypes with methylation status and thus computed eQTM (methylation to gene expression correlations) p-values, again retaining only the most significant eQTM for a gene.

## Model structure and learning

### Probabilistic graphical model

The pGENMi plate diagram is shown in Figure 1, which models the association between a specific TF and a phenotype – in this application drug response. There are four variable types in the plate diagram *(p,Zg,r_gm_,H)* and two parameter types (*a,w*_*m*_). The variable *r_gm_* is a binary indicator variable representing whether a gene *g* has a particular kind of evidence (e.g., cis-eQTL, cis-eQTM, etc.) of regulation by the TF. A rectangle labeled with an M engulfs r, indicating that there areM such kinds of regulatory evidence for each geneg: *r_gl_,r_g2_*,…*.,r_gM_.* Each *r_gm_* variable in a plate connects to the same *z_g_* variable. This variable is binary and latent (unshaded), and indicates whether expression of a gene *g* is related to the phenotype. A rectangle labeled with a *G* encapsulates *z_g_* and *r_gm_,* indicating that there is a latent variable *z_g_* and regulatory evidence vector *r_gm_* for each of *G* genes. Since *z_g_* is latent, its state is determined probabilistically, as shown below:

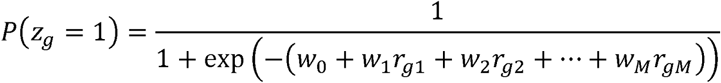

Here, the various evidences *r_gm_* for the regulation of gene *g* by the TF are combined in a logistic framework, each weighted by a coefficient *w_m_.* In this manner, a weighting of separate regulatory evidences determines whether a gene *g* is related to the phenotype. The variable p is a continuous variable in the range [0,1], representing the observed p-value of the Pearson correlation of a gene’s expression with a phenotype of interest. If a gene is related to the phenotype *(z_g_ =* 1), then we model p to follow a *Beta(α,* 1) distribution biased towards significant (small) p-values; *a* in this context is a shape parameter indicating the strength of the bias. It is also constrained to the range [0,1], with *α =* 1 equivalent to a uniform distribution over p-values. However, if a gene is unrelated to phenotype, we expect its expression-phenotype correlation p-value to be randomly distributed over the unit interval. These modeling assumptions are summarized below:

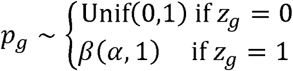

The binary variable *H* indicates the hypothesis to be tested. When *H =* 1, the model tries to find the best assignments to w and *α.* Traditionally, this is called the alternative hypothesis *or H_1_.* When *H =* 0, the model learns a smaller set of parameters, namely w_0_ and *a,* and the weights *w_1_*…*w_M_* are removed from the model entirely. In this case, the model explores how well it can explain the observed p-values without any regulatory evidences at all. Thus, *H =* 0 indicates the null hypothesis, or *H_0_.* We derive a score for how much better the alternative hypothesis explains the datay:

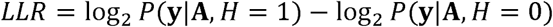

The pGENMi method uses the log likelihood ratio (LLR), computed for each TF separately, to rank TFs by their predicted association with the phenotype.

### Parameter estimation

The model finds assignments to the parameters, **w** = [*w_0_, w_t_*…*. w_M_]* and α, such that the likelihood of the observed p-values, ***p*** = [*p_a_*….*p_G_],* is maximized. We use the Expectation Maximization algorithm to find the optimal values for **w** and α. See **Supplementary Text 4** for more details. The optimization imposes no constraints on the parameters in the objective function. Thus, for instance, the weights of regulatory evidences (w) may be trained to negative values and thus reward genes that have regulatory evidence but no expression-phenotype correlation. As this is not consistent with our goals, pGENMi invokes a post-processing step to disregard the spurious TF-phenotype associations arising in such scenarios.

### Code Availability

Source code for the probabilistic graphical model will be available upon publication at the following location: http://veda.cs.uiuc.edu/pGENMi/

## Experimental validation data, methodology, and statistical analysis

### Cell culture and treatments

Human triple negative breast cancer (MDA-MB231), leukemia (Jurkat), and glioma (U251) cell lines were obtained from the American Type Culture Collection (Manassas, VA). MDA-MB-231 cells were cultured in L-15 medium containing 10% Fetal Bovine Serum (FBS). Jurkat cells were cultured in Roswell Park Memorial Institute (RPMI) 1640 Medium, containing 10% FBS. U251 cells were cultured in Dulbecco’s Modified Eagle Medium (DMEM), also containing 10% FBS.

The following 10 drugs were purchased from Sigma-Aldrich (St. Louis, MO): 6-Mercaptopurine (6-MP), 6-Thioguanine (6-TG), carboplatin, cisplatin (CDDP), cytarabine (Ara-C), docetaxel, epirubicin, gemcitabine, oxaliplatin, and paclitaxel. The remaining three drugs were obtained Selleck Chemicals (Houston, TX): cladribine, rapamycin (sirolimus), and temozolomide (TMZ). All 13 drugs were dissolved in dimethyl sulfoxide (DMSO) and aliquots of stock solutions were frozen at −80°C.

*RNA interference and Real-time quantitative reverse transcription-PCR (qRT-PCR)* siRNAs for candidate TFs and negative control siRNA were purchased from Dharmacon. Reverse transfection was performed for MDA-MB231 and U251 cells in 96-well plates. Specifically, 3000–4000 cells were mixed with 0.1 mL of lipofectamine RNAi-MAX reagent (Invitrogen) and 10 nM siRNA for each experiment. Electroporation was performed for Jurkat cells using Nucleofector^®^ Kit V from Lonza (Cologne, Germany).

Prior to electroporation, cells were washed with phosphate buffer saline (PBS) and counted. One million Jurkat cells were re-suspended in 100 µL of the Nucleofector^®^ Solution buffer and mixed with 100 nM of specific siRNA. The re-suspended cells were transferred to cuvettes and immediately electroporated using the program X-005. After electroporation, cells were incubated in a cuvette at room temperature for 10 minutes and then 500 µL of pre-warmed culture medium were added to the cuvette. Cells were then transferred to a 12-well plate and incubated at 37°C/5% CO_2_ overnight.

Total RNA was isolated from cultured cells transfected with control or TF-specific siRNAs with the Qiagen RNeasy kit (QIAGEN, Inc.), followed by qRT-PCR performed with the one-step Brilliant SYBR Green qRT-PCR master mix kit (Stratagene). Specifically, primers purchased from QIAGEN were used to perform qRT-PCR using the Stratagene Mx3005P Real-Time PCR detection system (Stratagene). All experiments were performed in triplicate with beta-actin as an internal control. Reverse transcribed Universal Human reference RNA (Stratagene) was used to generate a standard curve. Control reactions lacked RNA template.

### MTS cytotoxicity assay

Cell proliferation assays were performed in triplicates at each drug concentration. Cytotoxicity assays with the lymphoblastoid and tumor cell lines were performed in triplicates at each dose. Specifically, 90 μL of cells (5 × 10^4^ cells) were plated into 96-well plates (Corning, NY) and were treated with increasing does of a specific drug or radiation. After incubation for 72 hours, 20 μL of CellTiter 96^*®*^ AQueous Non-Radioactive Cell Proliferation Assay solution (Promega Corporation, Madison, WI) was added to each well. Plates were read in a Safire2 plate reader (Tecan AG, Switzerland).

Cytotoxicity assays with the tumor cell lines were performed using the CellTiter 96® AQueous Non-Radioactive Cell Proliferation Assay (Promega Corporation, Madison, WI). Specifically, 90 μL of cells (5 × 10^3^ cells) were plated into 96-well plates and were treated with increasing does of a specific drug. The escalation of concentrations for each drug is listed in **Supplementary Table 3**. After incubation for 72 hours, 20 μL of CellTiter 96® AQueous Non-Radioactive Cell Proliferation Assay solution (Promega Corporation, Madison, WI) was added to each well. Plates were read in a Safire2 plate reader (Tecan AG, Switzerland). Cytotoxicity was assessed by plotting cell survival versus drug concentration, on a log scale.

Radiation cytotoxicity was performed in triplicates at each radiation dose as described above. 100 μL of cells (5 × 10^3^ cells) were plated into 96-well plates and were treated with ionizing radiation at 0, 0.25, 0.5, 1, 2.5, 5, 10, and 20 Gy, using cesium-137 gamma-rays (J.L. Shepherd and Associates Mark I Model 25 Irradiator). After incubation for 72 hours, 20 μL of CellTiter 96 AQueous Non-Radioactive Cell Proliferation Assay solution was added to each well. Plates were read in a Safire2 plate reader (Tecan AG).

### Statistical analysis

Significance of the IC50 values between negative control siRNA and TF-specific siRNA was determined using a two-tailed unpaired t-test.

### Data Availability

Genotyped SNP and gene expression data are available in the NCBI Gene Expression Omnibus (GEO) (http://www.ncbi.nlm.nih.gov/geo) under SuperSeries accession no. GSE24277 and were originally published in a study by Niu et. al (55). In addition to gene expression and genotype data, the 444,797 methylation marks, originally published by Heyn et al. (56), were made available in the NCBI GEO under SuperSeries accession no. GSE36369. The cytotoxicity agents were all analyzed in a previous study (19). The cytotoxicity data that support the findings of this study are available from the corresponding author upon request.

## Acknowledgments

Funding for this work was provided in part by the NIH (Grant U54 GM114838 to SS, U19 GM61388 Pharmacogenomics Research Network and R01 CA138461 to LW) and in part by the Mayo Clinic-UIUC Alliance and by Grant 1U54GM114838 awarded by NIGMS through funds provided by the trans-NIH Big Data to Knowledge (BD2 K) initiative (www.bd2k.nih.gov). The content is solely the responsibility of the authors and does not necessarily represent the official views of the National Institutes of Health.

